# Design of a Cereblon construct for crystallographic and biophysical studies of protein degraders

**DOI:** 10.1101/2024.01.17.575503

**Authors:** Alena Kroupova, Valentina A. Spiteri, Hirotake Furihata, Darren Darren, Sarath Ramachandran, Zoe J. Rutter, Sohini Chakraborti, Kevin Haubrich, Julie Pethe, Denzel Gonzales, Andre Wijaya, Maria Rodriguez-Rios, Dylan M. Lynch, William Farnaby, Mark A. Nakasone, David Zollman, Alessio Ciulli

## Abstract

The ubiquitin E3 ligase cereblon (CRBN) is the target of therapeutic drugs thalidomide and lenalidomide and is recruited by most targeted protein degraders (PROTACs and molecular glues) in clinical development. Biophysical and structural investigation of CRBN has been limited by current constructs that either require co-expression with the adaptor DDB1 or inadequately represent full-length protein, with high-resolution structures of degraders ternary complexes remaining rare. We present the design of CRBN^midi^, a construct that readily expresses from *E. coli* with high yields as soluble, stable protein without DDB1. We benchmark CRBN^midi^ for wild-type functionality through a suite of biophysical techniques and solve high-resolution co-crystal structures of its binary and ternary complexes with degraders. We qualify CRBN^midi^ as an enabling tool to accelerate structure-based discovery of the next generation of CRBN based therapeutics.

**One sentence summary:** A novel Cereblon construct (CRBN^midi^) allows structural and biophysical enablement of ligand and degrader design

## Introduction

Targeted protein degradation enables development of small molecule therapies that promise to address medical needs that are currently unmet *via* conventional drugs (*1, 2*). Small-molecule degraders co-opt the ubiquitin-proteasome system by recruiting a specific target protein to a ubiquitin E3 ligase complex, to induce protein ubiquitination and subsequent degradation. This is achieved through bifunctional proteolysis-targeting chimeras (PROTACs) that are composed of two distinct portions that separately bind the target and E3 ligase, or so-called “molecular glues” that are typically smaller than PROTACs and bind preferentially or exclusively either the E3 ligase or the target (*3, 4*). To date, over 40 degraders between PROTACs and molecular glues, are under development in clinical trials for various disease conditions through targeting a range of proteins for degradation (*5, 6*).

Most of the current clinical candidates, and indeed many of the degraders reported in the literature to date, co-opt the protein cereblon (CRBN), the substrate receptor subunit of the Cullin4 E3 ligase complex (CRL4^CRBN^). CRBN was identified in 2010 as the target of the infamous drug thalidomide, and related immunomodulatory drugs (IMiDs) such as lenalidomide that is approved for treatment in multiple myeloma and other haematological malignancies (*7, 8*). Thalidomide and lenalidomide bind non-covalently to CRBN with structurally defined binding modes that mimic the C-terminal cyclic imide degron of CRBN substrates (*9-11*). Upon binding to CRBN, IMiDs act as molecular glues, enhancing the recruitment of a wide range of zinc-finger containing transcription factors and other proteins as CRBN *neo*-substrates leading to their rapid ubiquitination and degradation (*12-14*). The degradation activity of IMiDs underpins their potent pleiotropic activity and explained thalidomide teratogenicity and lenalidomide anti-tumour efficacy. Thalidomide, lenalidomide and other CRBN binders or related chemotypes provide attractive drug-like starting points to medicinal chemistry development of glue-type CRBN modulators as well as binding moieties for the design of PROTAC degraders co-opting CRBN as the E3 ligase (*15*).

Despite the progress to date, the development of CRBN-based molecular glue and PROTAC degraders has witnessed limited enablement through structure-guided drug design. Indeed, only few high-resolution structures of ternary complexes CRBN:degrader:target have been solved using either X-ray crystallography or cryo-EM (*16-24*). Similarly, *in vitro* biophysical characterization of CRBN-based degraders has remained sparse (*24-27*). This is in striking contrast with the structural and biophysical enablement that supports design of PROTAC degraders recruiting the other popular E3 ligase, von Hippel Lindau (VHL), as illustrated by the many ternary complex co-crystal structures solved by us and others that have led to efficient structure-guided design of VHL-based PROTACs (*28-34*), and their biophysical characterization (*28, 35, 36*). Ternary co-crystal structures have also enabled structure-driven design of Cyclin K degraders which glue CDK12 to the CRL4 adaptor protein DDB1 (Damage specific DNA binding protein 1), to induce ubiquitination and degradation of CDK12-complexed Cyclin K (*37-40*). We therefore hypothesized that full structural enablement of CRBN could substantially accelerate the design of CRBN-based degraders, and so turned our attention to protein construct design.

### Design of CRBN^midi^: a truncated CRBN construct containing Lon and TBD domains

We hypothesized that a major gap to structural enablement of CRBN has been largely caused by the challenges in producing and handling a suitable recombinant CRBN protein construct. CRBN consists of an unstructured N-terminal region and three folded domains: the Lon protease-like domain (Lon), the helical bundle (HB) which facilitates binding to the adaptor protein DDB1, and the thalidomide binding domain (TBD) (Fig. 1A). Upon ligand binding, the TBD and Lon domains undergo a ∼45° rearrangement between an apo ‘open’ and ligand-bound ‘closed’ state (*16, 19, 23*). Recombinant CRBN is insoluble when expressed in isolation but can be purified in complex with DDB1 from insect cells (*9*). Whilst suitable for *in vitro* biochemical characterization, this full-length construct has not been crystallized so far. Several approaches have been developed to enable crystallization, including (i) a chimeric complex of *G. gallus* CRBN orthologue with human DDB1 (CRBN^*G*.*gallus*^:DDB1^*H*.*sapiens*^) (*9*); (ii) a construct lacking the unstructured N-terminal region of CRBN (CRBN^ΔN^:DDB1) (*17, 20, 22, 41*); and (iii) DDB1 lacking the WD40 propeller B (CRBN^ΔN^:DDB1^ΔBPB^)(*16, 18, 19, 21, 24*). The majority of the crystal structures that have resulted from these constructs have been solved to a resolution lower than 3 Å. Similarly, the full-length CRBN:DDB1 complex has been recently shown to be amenable to cryo-EM studies, however with overall resolution below 3 Å and often poorly resolved maps of the CRBN ligand binding site (*23*). Alternatively, the isolated TBD of CRBN, spanning residues 328-426 (CRBN^TBD^), has been widely used because it is stable in solution and expressible in large quantities from *E. coli*, with either the human sequence (*42*), its murine orthologue (*28, 30*) or the single-domain orthologue *M. gryphiswaldense* CRBN isoform 4 (*43, 44*), which all crystallize routinely to sub-2 Å resolution. However, the TBD domain alone lacks a significant portion of the full-length CRBN protein and therefore is not representative for studies of ternary complexes which could involve interactions with the missing Lon domain.

**Fig. 1.**
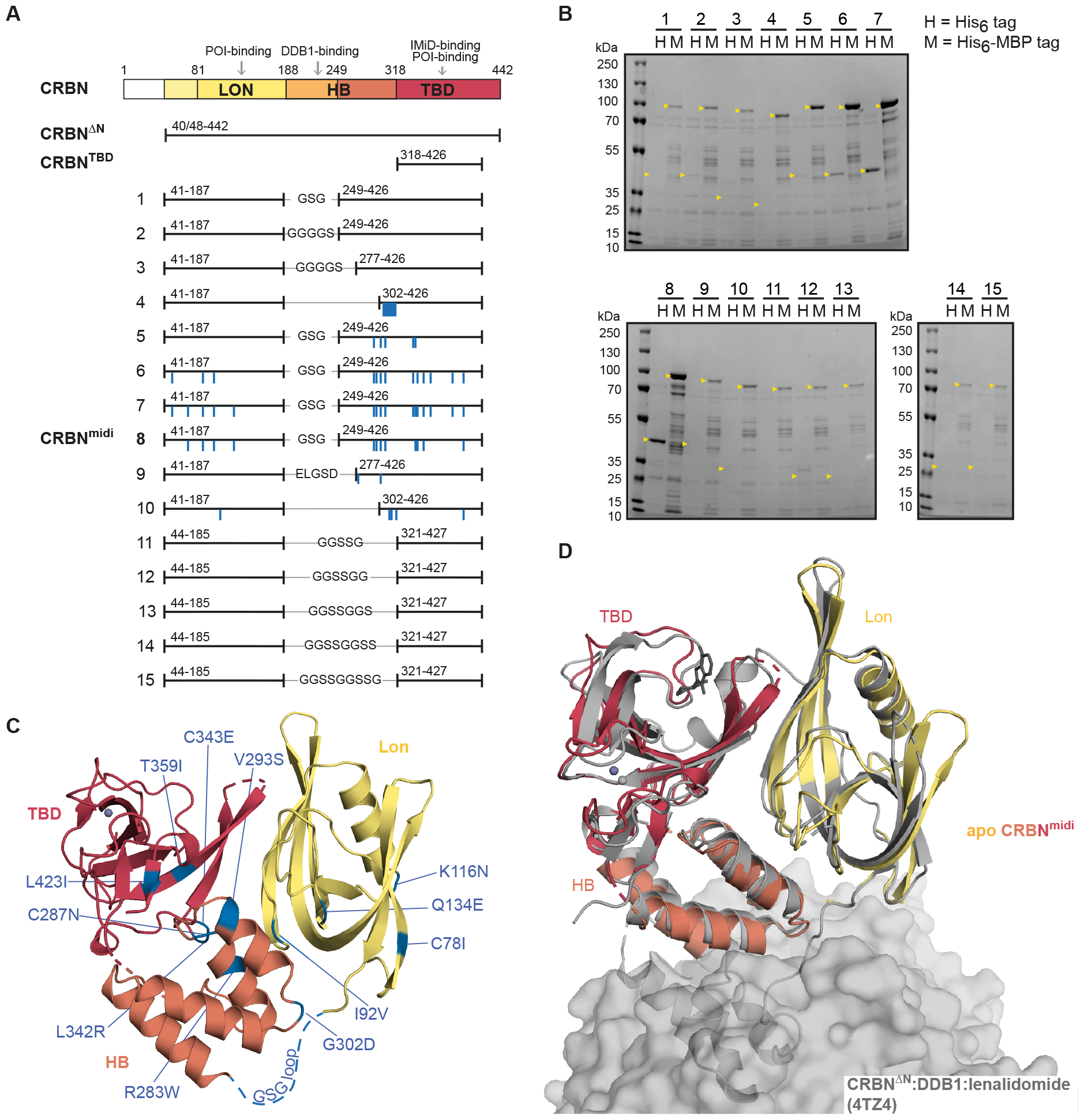
Design and characterization of CRBN^midi^. (**A**) Domain composition of full-length CRBN (top), constructs previously used for structural and biophysical characterization, CRBN^ΔN^ and CRBN^TBD^, and constructs **1**-**15** designed in this study. The most promising construct 8 was designated as CRBN^midi^. Mutated residues are indicated as blue lines. (**B**) Ni-NTA affinity pull-down assay of lysates from *E. coli* BL21(DE3) cells expressing candidate constructs **1**-**15** with an N-terminal His_6_ (H) or His_6_-MBP (M) tag. The expected size of each construct is marked with a yellow arrow. (**C**) Crystal structure of the apo CRBN^midi^ containing Lon (yellow), HB (orange) and TBD (red) domains. The mutated residues are indicated (blue), the unresolved region containing the GSG linker is shown as blue dashed line, Zn^2+^ is shown as a purple sphere. (**D**) Superposition of apo CRBN^midi^ (yellow, orange, red) with CRBN^ΔN^:DDB1 bound to lenalidomide (grey, PDB ID: 4TZ4). DDB1 is shown as a grey surface.

To address these limitations, we set out to develop a truncated CRBN construct containing both TBD and Lon domains, that would conveniently express soluble and stable protein from *E. coli* with comparable functionality to the full-length wild-type CRBN. To this end, we rationally designed fifteen new constructs using three main strategies: (i) partial deletions of the HB domain (constructs **1**-**4**), (ii) introducing stabilizing mutations in a construct lacking the DDB1-interacting region (constructs **5**-**10**), and (iii) deletion of the full HB domain (constructs **11**-**15**) (Fig. 1A, Table 1). In detail, firstly we removed the DDB1-binding region of the HB domain, containing residues 188-248, which in absence of DDB1 exposes a hydrophobic patch prone to inducing aggregation of CRBN, resulting in constructs **1** and **2** with GSG or GGGGS linkers, respectively. Further truncations of the remaining HB domain were guided by secondary structure boundaries. Construct **3** contains two C-terminal alpha-helices from the HB domain spanning residues 277-317 with a GGGGS linker and construct **4** spans HB residues 302-317, where Ile305-Lys317 was mutated to a polyAla helix. The removal of portions of the HB domain resulted in previously buried residues becoming solvent exposed, in some cases compromising the solubility of the construct. In our second approach, we used the Protein Repair One-Stop Shop (PROSS) server (*45*) to identify amino acid substitutions that would positively contribute to protein stability and solubility. Constructs **5**-**10** were designed by manually curating the results of the PROSS analysis to exclude any residues within, and in the vicinity of, the IMiD-binding site that would engage in protein-protein interactions during ternary complex formation (*16-24*). Thirdly, the HB domain was completely removed in constructs **11**-**15** to contain only the TBD and Lon domains connected by an assortment of flexible linkers and the C366S mutation previously introduced to improve crystallization of the TBD domain (*42*). In all constructs, glycine and serine-rich linkers were chosen for their flexibility (*46*) with the length requirement predicted for each truncation by analysis of published CRBN crystal structures.

**Table 1.** Design of CRBN constructs.

| Construct ID | N-terminal domain | Linker | C-terminal domain | Mutations |
| --- | --- | --- | --- | --- |
| 1 | 41-187 | GSG | 249-426 | - |
| 2 | 41-187 | GGGGS | 249-426 | - |
| 3 | 41-187 | GGGGS | 277-426 | - |
| 4 | 41-187 | GGGGS | 302-426 | I305-K317 -> polyA |
| 5 | 41-187 | GSG | 249-426 | R283W, V293S, G302D, L340M, C343E |
| 6 | 41-187 | GSG | 249-426 | T58S, I92V, K116N, R283W, C287N, V293S, G302D, L340M, C343E, T359I, C366K, S410R, L423I |
| 7 | 41-187 | GSG | 249-426 | T58S, C78I, I92V, K116N, Q134E, R283W, C287N, V293S, G302D, L340M, L342R, C343E, T359I, C366K, S410R, L423I |
| 8 | 41-187 | GSG | 249-426 | C78I, I92V, K116N, Q134E, R283W, C287N, V293S, G302D, L342R, C343E, T359I, L423I |
| 9 | 41-187 | ELGSD | 277-426 | P277T, V293H |
| 10 | 41-187 | - | 302-426 | V128R, A304K, Q306D, C310H, L423Q |
| 11 | 44-185 | GGSSG | 321-427 | C366S |
| 12 | 44-185 | GGSSGG | 321-427 | C366S |
| 13 | 44-185 | GGSSGGS | 321-427 | C366S |
| 14 | 44-185 | GGSSGGSS | 321-427 | C366S |
| 15 | 44-185 | GGSSGGSSG | 321-427 | C366S |

To assess protein expression and solubility, we cloned all constructs with an N-terminal His_6_ or His_6_ coupled to maltose binding protein (MBP) affinity tags and performed Ni-NTA affinity pull-down assay from lysates of cells transformed with each construct (Fig. 1B). All His_6_-MBP-tagged constructs showed detectable immobilized protein levels whereas only constructs **2** and **5**-**8** immobilized above the detection limit in the absence of the MBP tag. Intriguingly, the stabilizing mutations introduced based on PROSS analysis proved to have important effects, with the best expressing construct (construct **8**, hereafter referred to as CRBN^midi^) incorporating 12 mutations. CRBN^midi^ protein was subsequently expressed and purified in large scale to assess its yield and homogeneity. The purification involved Ni-NTA affinity chromatography, desalting, tag cleavage, and reverse Ni-NTA affinity chromatography followed by size-exclusion chromatography (SEC) which yielded a pure monodisperse sample as confirmed by SDS-PAGE and SEC (Fig. S1) resulting in 1.5 mg of purified protein per 1 L of culture. Next, we successfully crystallized CRBN^midi^ protein and determined its crystal structure to a resolution of 3.11 Å (Fig. 1C, Table S1). The fold of CRBN^midi^ superposes well with the corresponding portion of CRBN^ΔN^:DDB1:lenalidomide (PDB ID: 4TZ4) with a RMSD of 1.27 Å (over 232 out of 275 Cα atoms) indicating the new construct adopts a biologically relevant fold equivalent to that of the wild-type protein. As with CRBN^ΔN^:DDB1, the structure shows CRBN^midi^ co-purified with a Zn^2+^ ion bound to the zinc finger of the TBD. Despite the absence of a small-molecule ligand, CRBN^midi^ adopted the “closed conformation” within the crystal.

### CRBN^midi^ binary complexes reveal molecular insights into ligand recognition

We next assessed the suitability of CRBN^midi^ for accessing protein-ligand binary complexes. We determined the co-crystal structures of CRBN^midi^ in complex with mezigdomide or lenalidomide to a resolution of 2.19 and 2.50 Å, respectively (Table S1). The CRBN^midi^:mezigdomide structure revealed a binding mode consistent with a previously published CRBN:DDB1:mezigdomide cryo-EM structure (PDB ID: 8D7U) with superposition yielding RMSD of 0.67 Å (over 259 out of 306 Cα atoms) (Fig. 2A). In addition, the crystal structure of mezigdomide-bound CRBN^midi^ revealed a non-covalent interaction between the fluorine atom of the ligand and Asp149 which was previously undetected in the CRBN:DDB1:mezigdomide cryo-EM structure (*23*) due to lower map quality around the ligand binding site (Fig. 2A middle panel). As mezigdomide interacts with both the TBD and Lon domains of CRBN, the CRBN^midi^:mezigdomide structure illustrates the importance of studying binary interactions using a construct containing both TBD and Lon domains in contrast to the CRBN^TBD^ construct often used for binary studies. Similarly to the CRBN^midi^:mezigdomide structure, the crystal structure of CRBN^midi^ bound to lenalidomide shows the closed conformation (Fig 2B) and superposition with CRBN^ΔN^:DDB1:lenalidomide (PDB ID: 4TZ4, RMSD of 1.22 Å over 227 out of 288 Cα atoms) indicates conserved ligand binding. Together, these high-resolution structures of CRBN^midi^ with bound ligands of different affinities establish it as an enabling construct for X-ray crystallographic structure-based design.

**Fig. 2.**
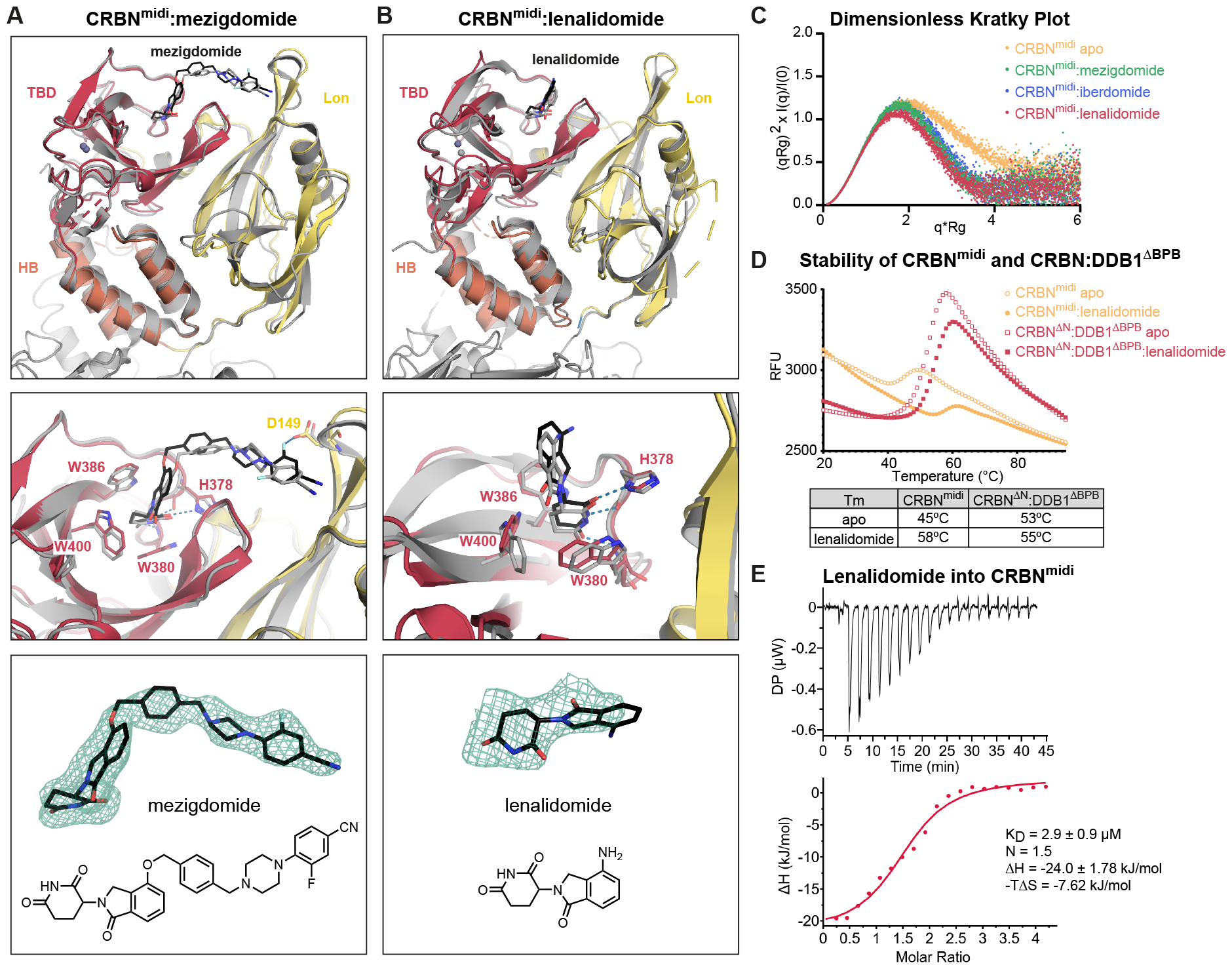
Structural and biophysical analysis of binary protein-ligand complexes using CRBN^midi^. (**A**) Crystal structure of CRBN^midi^ (domains coloured as in Fig. 1) bound to mezigdomide (shown as black sticks) superposed with the cryo-EM structure of CRBN:DDB1:mezigdomide (PDB ID: 8D7U). (**B**) Crystal structure of CRBN^midi^ (domains coloured as in Fig. 1) bound to lenalidomide (shown as black sticks) superposed with crystal structure of CRBN^ΔN^:DDB1:lenalidomide (grey, PDB ID: 4TZ4). (**A-B**) Overall fold shown as cartoon (top); detail of the ligand binding site (middle) with residues involved in ligand binding shown as sticks and potential H-bonds shown as blue dashed lines; Polder OMIT (Fo-Fc) map of the ligands contoured to 3 σ shown as green mesh, alongside the chemical structures of the compounds (bottom panel). (**C**) Dimensionless Kratky Plot generated from SAXS data of apo CRBN^midi^ (yellow) and CRBN^midi^ bound to mezigdomide (green), iberdomide (blue), or lenalidomide (red). (**D**) Differential scanning fluorimetry (DSF) denaturing curves for CRBN^midi^ (yellow) and CRBN^ΔN^:DDB1^ΔBPB^ (red) in the absence (empty circles or squares, respectively) or presence of lenalidomide (filled circles or squares). (**E**) ITC measurement of lenalidomide binding to CRBN^midi^.

CRBN transitions from an open to a closed conformation upon ligand binding, with major hinge bending between the TBD and Lon domains (*16, 19, 23*). Our extensive engineering of the linker region between these two domains in CRBN^midi^ prompted us to next conduct biophysical experiments to compare the behaviour of CRBN^midi^ and full-length wild-type CRBN in solution. Whilst both apo and ligand-bound CRBN^midi^ crystallized in the closed conformation, we used small-angle X-ray scattering (SAXS) coupled to in-line SEC to analyse the domain rearrangement of CRBN^midi^ in solution (Fig. 2C, S2). In the absence of a binder, the radius of gyration (R_G_) for CRBN^midi^ was 26.73 Å, corresponding well to the predicted R_G_ of 25.65 Å for apo CRBN^midi^ in its open conformation, as modelled based on the open conformation of full-length CRBN cryo-EM structure (PDB ID: 8CVP)(*23*). A theoretical scattering curve for the model of the open conformation fits the experimental curve well (Fig. S2). In the presence of ligand, we observed a R_G_ of 23.69, 22.67 and 22.49 Å for iberdomide, mezigdomide and lenalidomide-bound CRBN^midi^, respectively, which were significantly smaller than in the absence of ligand (26.73 Å). Experimental and theoretical scattering curves and R_G_ correlated well for liganded CRBN^midi^ as well (c.f. 22.67 Å and 23.76 Å, respectively, for the complex with mezigdomide, Fig. S2). To compare the extent of flexibility of the apo and ligand-bound protein we used the dimensionless Kratky plot (*47*), a representation of the SAXS data normalised for scattering intensity and R_G_ (Fig. 2C). It indicates that apo CRBN^midi^ shows significant flexibility and can access a larger conformational landscape, evidenced by the tail of the bell curve trailing higher than that of the ligand-bound CRBN^midi^ curves. In contrast, the curves for the ligand-bound return to the baseline and show a peak maximum close to the Guinier-Kratky point, indicating a rigid, globular conformation. This is consistent with the findings from Watson *et al*. that show that apo cereblon can readily access open conformations in cryo-EM studies (*23*). Furthermore, molecular dynamics (MD) simulation on our CRBN^midi^:mezigdomide crystal structure alongside simulation of the corresponding cryo-EM structure (PDB ID: 8D7U)(*23*) was next performed to computationally investigate and compare the dynamics of the protein residues and the interaction profile of the ligand in both protein structures. The Root Mean Square Fluctuations (RMSF) of the residues and protein-ligand interaction profiles as observed from the MD simulations indicated that CRBN^midi^ behaves similarly to the WT construct (Fig S3, Tables S4-S6, see Supplementary text for details). These data evidence overall compaction of CRBN^midi^ protein undergoing the open-close conformational rearrangement upon ligand binding, in a manner consistent with wild-type full-length CRBN, and exemplify the sensitivity of SAXS for monitoring binding of CRBN ligands which can modulate this open-close equilibrium.

To gain further insights into the biophysical thermodynamics of the protein-ligand binding equilibria, we performed differential scanning fluorimetry (DSF) and isothermal titration calorimetry (ITC) studies with CRBN^midi^. Using DSF, we compared the thermal denaturation curves of lenalidomide-bound and unbound CRBN^midi^ with those of CRBN^ΔN^:DDB1^ΔBPB^ (Fig. 2D). CRBN^midi^ had a melting temperature (T_m_) of 45 °C that was remarkably stabilized to 58 °C in the presence of lenalidomide (leading to a large ΔT_m_ of 13°C). In comparison, CRBN^ΔN^:DDB1^ΔBPB^ showed a T_m_ of 53°C in the absence of ligand and T_m_ of 55°C in the presence of lenalidomide (ΔT_m_ = 2°C). This much larger ΔT_m_ upon ligand binding qualifies CRBN^midi^ as a suitable construct for high-throughput screening of potential CRBN binders and modulators by DSF. Next, we obtained binary binding affinity measurements by ITC. Titration of lenalidomide into CRBN^midi^ resulted in a conventional sigmoidal curve with endothermic profile, and data fitting yielded a K_D_ of 2.9 μM and H of –24 kJ/mol (Fig. 2E). For reference, ITC titration data have previously been reported for lenalidomide against CRBN^TBD^ (K_D_ of 19 μM, H of –21.8 kJ/mol) and CRBN:DDB1 (K_D_ of 0.6 μM, H of 10.5 kJ/mol) (*48*). The differences in binding affinity and enthalpy observed are consistent with the use of different protein constructs and with differences in experimental conditions, meaning absolute K_D_ and H values are not directly comparable. Nonetheless, our data benchmarks CRBN^midi^ against widely used current constructs and qualifies CRBN^midi^ as relevant and suitable for ITC analysis of protein-ligand binding. Overall, we show that CRBN^midi^ enables in-depth structural and biophysical analysis of binary interactions *via* a wide range of biophysical and structural techniques thus qualifying it as suitable to library screening for identification and characterization of new binders.

### High-resolution crystal structures of degrader ternary complexes enabled by CRBN^midi^

We set out to exemplify the utility of CRBN^midi^ for enabling co-crystal structures of ternary complexes formed by molecular glue or PROTAC degraders. We first determined the crystal structure of CRBN^midi^ in complex with mezigdomide and the second zinc-finger of neo-substrate Ikaros (IKZF1^ZF2^) to a resolution of 2.15 Å (Fig. 3A-C, Table S2). Superposition of our new X-ray structure with the cryo-EM structure of CRBN:DDB1:mezigdomide:IKZF1^ZF1-2-3^ (PDB ID: 8D7Z) (*23*) (RMSD of 0.85 Å over 245 out of 293 Cα atoms) shows a highly consistent binding mode of both IKZF1 and mezigdomide engaged in the ternary complex with CRBN (Fig. 3B). Importantly, the higher resolution of our crystal structure and greater quality of the density map at the binding interface allowed us to unambiguously observe several protein-ligand and protein-protein interactions, including contacts that had not previously been identified (Fig. 3C). An extended network of mezigdomide-CRBN contacts is observed. At the phthalimide-end, the ligand’s glutarimide moiety interacts *via* the canonical three hydrogen bonds with His378^CRBN^ and Trp380^CRBN^, while the oxoisoindoline carbonyl group forms a hydrogen bond with the side-chain NH_2_ of Asn351^CRBN^. Mezigdomide adopts a characteristic U-like shape around its central phenyl ring, wrapped around Pro352 and His353 at the tip of the beta-hairpin sensor loop (residues 341-361), conformation stabilized through several π-π stacks and hydrophobic interactions (Fig. 3C). At the other end, the piperidine-benzonitrile moiety of mezigdomide fits snugly into a hydrophobic pocket defined on one side by Phe102, Phe150 and Ile152 from the CRBN’s Lon domain, and on the other side by the tip of the sensor loop (Fig. 3C). In contrast to the extensive contacts between mezigdomide and CRBN, no further interactions are observed between mezigdomide and IKZF1^ZF2^ beyond the known stacking of the neo-substrate Gly-loop on top of the isoindolinone ring, consistent with the molecular glue degrader binding exclusively to CRBN and not IKFZ1 at binary level (*16, 19*). In addition, we observed several direct protein-protein contacts mediated by the degrader, also not all previously observed: Asn148^IKZF1^, *via* hydrogen bond from its backbone carbonyl to the side chain NH_2_ of Asn351^CRBN^, and through side-chain packing with His353^CRBN^; Gln149^IKZF1^, *via* hydrogen bond from its backbone carbonyl to the side chain NH_2_ of Asn351^CRBN^, and side-chain contacts with Tyr355^CRBN^; and Cys150^IKZF1^, *via* a hydrogen bond from the backbone carbonyl to the side chain NH of Trp400^CRBN^ (Fig. 3C). In our structure, Cys150^IKZF1^ is not within hydrogen bonding distance of His397^CRBN^, unlike the cryo-EM structure (*23*). Moreover, compared to our binary CRBN^midi^:mezigdomide structure (Fig. 2A), the fluorine of mezigdomide does not engage Asp149 in the ternary structure with IKZF1^ZF2^. Overall, our co-crystal structure has allowed us to detail many protein-ligand and protein-protein interactions within the ternary complex at an atomic resolution not previously achieved.

**Fig. 3.**
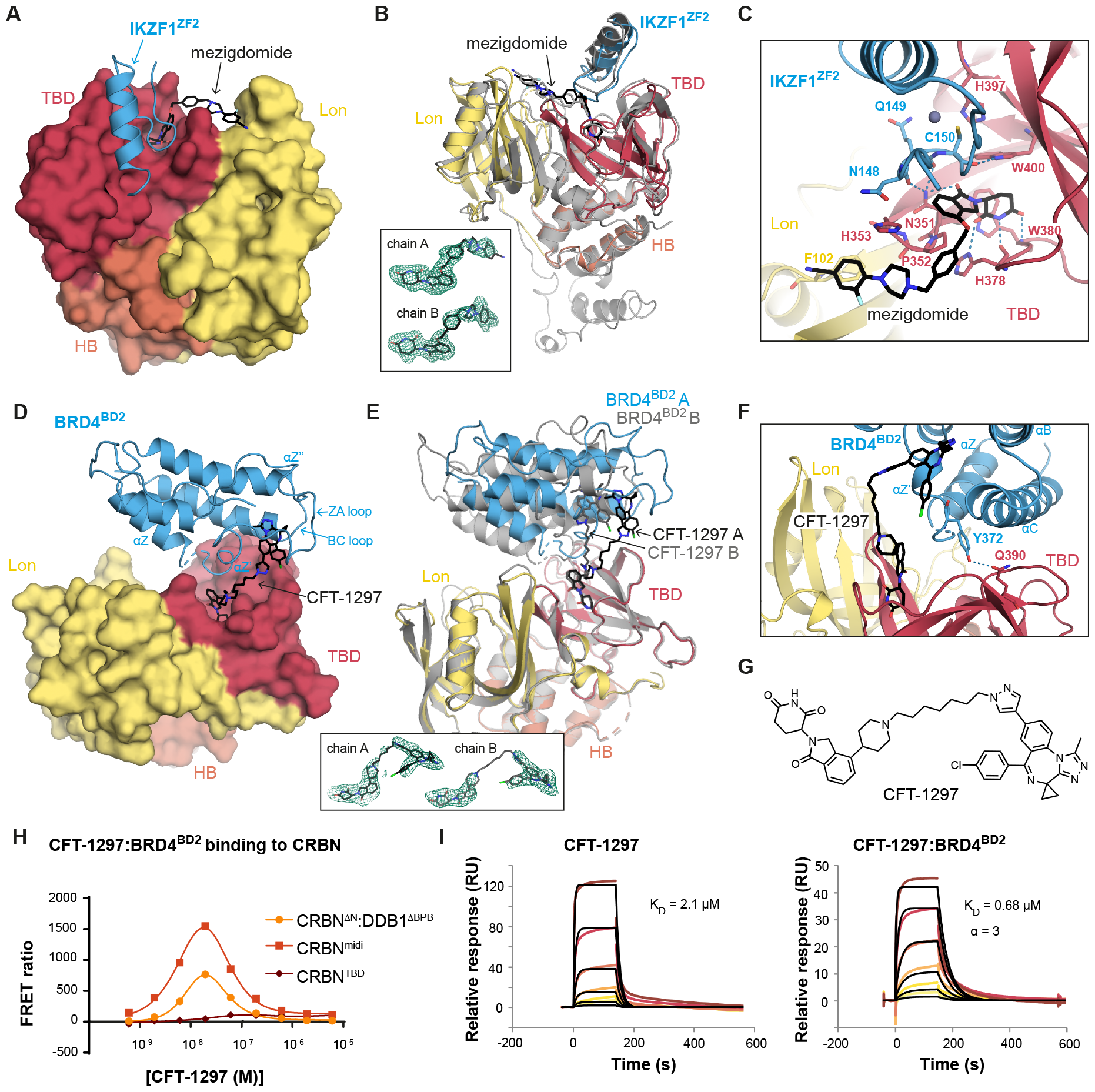
Ternary complex characterization using CRBN^midi^. (**A-C**) Crystal structure of CRBN^midi^ in complex with mezigdomide and IKZF1^ZF2^. (**A**) CRBN^midi^ is shown in surface representation, mezigdomide is shown as black sticks and IKZF1^ZF2^ is shown as blue cartoon. (**B**) Superposition of the CRBN^midi^:mezigdomide:IKZF1^ZF2^ crystal structure (coloured as in A) with the cryo-EM structure of CRBN:DDB1:mezigdomide:IKZF1^ZF1-2-3^ (PDB ID: 8D7Z, grey). Polder OMIT (Fo-Fc) map of mezigdomide in protomers A and B contoured to 3 σ is shown in the inset. (**C**) Close-up of the ligand-protein and protein-protein interaction network in (B). Hydrogen bonding interactions in the CRBN^midi^:mezigdomide:IKZF1^ZF2^ interface are shown as blue dashed lines. (**D-G**) Crystal structure of CRBN^midi^ in complex with CFT-1297 and BRD4^BD2^. (**D**) CRBN^midi^ is shown in surface representation, CFT-1297 is shown as black sticks and BRD4^BD2^ is shown as blue cartoon. (**E**) The two protomers in the asymmetric unit, chain A coloured, chain B in grey. Polder (OMIT) (Fo-Fc) map of CFT-1297 in chains A and B contoured to 3 σ shown in the inset. (**F**) Close-up of the interface between BRD4^BD2^ (chain B) and CRBN^midi^, residues Tyr372^BRD4^ and Gln390^CRBN^ are shown as sticks with the hydrogen bond shown as blue dashed line. (**G**) Chemical structure of CFT-1297. (**H**) TR-FRET traces monitoring ternary complex formation of BRD4^BD2^ and different constructs of CRBN in the presence of a dilution series of CFT-1297. (**I**) CRBN^midi^-on-chip SPR sensograms measuring binding of CFT-1297 alone (left) and CFT-1297 pre-incubated with BRD4^BD2^ (right).

We next solved the ternary crystal structure of CRBN^midi^ in complex with PROTAC CFT-1297 and BRD4^BD2^ to a resolution of 2.9 Å (Fig. 3D-G, Table S2). We chose compound CFT-1297 as a BET PROTAC degrader for crystallographic studies because it is reported to exhibit positive cooperativity of ternary complex formation, and was also further characterized by hydrogen-deuterium exchange mass spectrometry (HDX-MS) (*25*), providing some low-resolution structural information. The structure contains two protomers in the asymmetric unit which show a slightly different bound pose of the BRD4^BD2^ molecules relative to CRBN, caused by a tilt of 27° of the PROTAC around the piperidine moiety that connects the CRBN binder and the alkylic linker (Fig. 3E). The bromodomain docks sideways on top of the CRBN TBD, an orientation imposed by the long rigid alkylic linker of the PROTAC. The surface of the bromodomain most buried within the complex comprise the region from the end of the Z-helix to the middle of the ZA-loop, including the αZ’ helix from the ZA loop (residues 362-378). This region packs against the central linker region of the PROTAC at one end, and the CRBN TBD between the sensor loop and residues 390-396 at the other end, where a hydrogen bond is observed between the side chains of Tyr372^BRD4^ and Gln390^CRBN^ (Fig. 3F). The remainder of the ZA loop (residues 379-400) and the whole BC-loop are solvent-exposed within the ternary complex structure, as well as the Lon domain of CRBN which is also fully solvent-exposed (Fig. 3D). These observations are inconsistent with the HDX-MS results from Eron *et al*. where shielding of the BC loop and Lon domain in Brd4^BD1^ and CRBN, respectively, were observed in the ternary complex (*25*), suggesting a differential mode of recognition for BD1 *vs*. BD2. Additional protein-protein hydrophobic contacts are also observed as the C-helix of the bromodomain (residues 438-449) is found packed against CRBN at the opposite side of the TBD, with the side chains of Glu438^BRD4^ and Arg373^CRBN^ at this interface located at a potential salt-bridge distance. Overall, our novel co-crystal structure reveals at atomic resolution an extensive network of interactions mediating the recognition mode of a CRBN:PROTAC:Brd4 ternary complex, and suggest distinct ternary complex recruitment mode of the two bromodomains of Brd4 to CRBN by CFT-1297.

To establish the utility of CRBN^midi^ to biophysical studies of PROTAC ternary complexes in solution, we next performed TR-FRET and SPR binding assays. First, we monitored the proximity between BRD4^BD2^ and CRBN^midi^ upon increasing concentrations of CFT-1297 using TR-FRET, alongside CRBN^TBD^ and wild-type CRBN^ΔN^:DDB1^ΔBPB^ to allow benchmarking of our new construct (Fig. 3H). CRBN^ΔN^:DDB1^ΔBPB^ and CRBN^midi^ produced the expected bell-shaped profile exhibiting the characteristic hook effect at high concentration of bifunctional molecule, and with near identical maxima concentrations (17 nM vs. 21 nM, respectively) and comparable broadness of the curves. In contrast, no significant ternary complex formation was detected for the CRBN^TBD^ construct, highlighting the importance of using a CRBN construct that includes the Lon domain for biophysical characterization of degrader ternary complexes. With formation of the ternary complex CRBN^midi^:CFT-1297: BRD4^BD2^ confirmed by TR-FRET, we next aimed to quantitatively measure the binding affinity, cooperativity and kinetic stability of this species using an SPR ternary complex assay conceptually similar to the one previously developed by us with VHL (*35*) and by Bonazzi *et al*. and Ma *et al*. for CRBN:DDB1 (*24, 27*). First, we immobilised CRBN^midi^ on a Ni-NTA SPR chip *via* its His-tag and monitored the binding of CFT-1297 to CRBN^midi^ in the absence or presence of saturating amounts of BRD4^BD2^ (Fig. 3I, and Table S3). We observed a 3-fold enhanced affinity of CRBN for the PROTAC:BRD4^BD2^ binary complex when compared to that for the PROTAC alone (ternary K_D_ = 0.68 μM *vs*. binary K_D_ = 2.1 μM) and a corresponding increase in the stability of ternary *vs*. binary complex as measured from dissociation kinetics (t_1/2_ = 31 s *vs*. 9 s). This positive cooperativity (α = 3) and increased durability of CRBN E3 ligase engagement suggest favourability of the extensive network of intra-complex contacts observed in our ternary co-crystal structure.

## Conclusion

Targeted degradation has shown considerable promise as a route for therapeutic intervention. Co-opting of the E3 ligase CRBN is an established viable approach by using molecular glue or PROTAC degraders to productively engage with target proteins to trigger their rapid ubiquitination and degradation *in vivo*. The formation of the ternary complex is the key mechanistic step in the mechanism of action of small molecule degraders. It is established that characteristics such as affinity and half-life of the ternary complex correlate well with degradation fitness (*26, 29, 35, 36, 40*), and potentiating ternary complex affinity, stability and favourability to target ubiquitination provides a powerful optimization strategy. This goal can be expediated through biophysical and structural insights of CRBN recruitment, ushering structure-guided design of novel chemical matter. Our CRBN^midi^ construct provides a new reagent that extends and enables future endeavours in several ways. First, the nature of our engineering, including both TBD and Lon domains in a single construct, addresses and solves many limitations of current constructs, as illustrated by convenient high-level expression in *E. coli*, solubility, and stability as monomeric protein without the need to co-express with DDB1, as well as apparent readiness to crystallize in ligand-bound forms. Second, we exemplify the utility of CRBN^midi^ in allowing X-ray crystallography of CRBN ternary complexes that had not been crystallized before and at higher resolution than previously attained with other constructs through either X-ray or cryo-EM investigations (*9, 16-18, 20-25, 41*). Third, we demonstrate suitability of CRBN^midi^ for biophysical ligand binding studies in solution, and provide robust benchmark against wild-type functionality through a variety of assay read-outs and settings *via* techniques including SAXS, DSF, TR-FRET, ITC and SPR. The soluble, stable, and readily producible CRBN^midi^ that we here disclose and validate could be useful not only for enabling structure-based screens and designs for therapeutic applications but also as a facile molecular tool for probing native interactions and structure-function properties of CRBN.

Moving forward, there are many avenues to pursue leveraging our construct and the biophysical and structural assays described here and beyond to accelerate ligand screening and design, and to illuminate molecular mechanisms. The robustness and convenience of CRBN^midi^ and its proneness to yielding high-resolution liganded co-crystal structures as evidenced in this study open the door to a wide range of refined strategies for the design and optimization of protein degradation therapies. As with any new tools, the utility and applicability of CRBN^midi^ must be duly considered for each application, and there remain caveats. For example, to aid crystallizability, we have deleted the flexible 40 N-terminal residues that, albeit not essential, have been shown to contribute to stabilizing the closed conformation of CRBN (*23*). Similarly, whilst CRBN^midi^ is deemed acceptable to determine the immediate interactions within the ternary complexes, it will remain a surrogate for studying the native CRL4^CRBN^ E3 ligase complexes. Nonetheless, we anticipate that the data reported in this study and the open-access availability of the construct e.g. *via* Addgene, will aid broad uptake by the community and lead to many new advances in the targeted protein degradation field, ultimately accelerating the discovery of degrader therapeutics.

## Supporting information

Data S1 containing Tables S4 to S6

## Acknowledgements

We thank Adam Bond for the gift of BRD4^BD2^ protein, Diane Cassidy for technical support, and Moriz Mayer (Boehringer Ingelheim) for performing confirmatory NMR spectra with compound CFT-1297. We acknowledge the Diamond Light Source (proposals mx26793, mx35324, and sm33832) for provision of synchrotron radiation facilities, and we would like to thank staff of beamlines I04, I24, and B21 for assistance and support with crystal testing and data collection.

## Author contributions

Conceptualization: SR, DZ, AC. Methodology: AK, VAS, SR, SC, WF, DZ. Investigation: AK, VAS, HF, DD, SR, ZR, SC, KH, JP, DG, AW, MRR, DML, MN, WF, DZ. Visualization: AK, VAS, HF, SR, SC, KH, DZ. Funding acquisition: AC. Supervision: SR, WF, DZ, AC. Writing – original draft: AK, VAS, HF, DD, ZR, SC, KH, DZ. Writing – review & editing: AK, VAS, WF, DZ, AC

## Data and materials availability

Atomic models and structure factors have been deposited to the Protein Data Bank (PDB) under accession numbers: 8RQ1, 8RQ8, 8RQ9, 8RQA, 8RQC. The SAXS data have been deposited to Small Angle Scattering Biological Data Bank (SASBDB) and are awaiting approval. The His_6_-CRBN^midi^ expression plasmid is in the process of being deposited to Addgene.

