## Supplementary Materials for "Design of a Cereblon construct for crystallographic and biophysical studies of protein degraders"

**The PDF file includes:**

Materials and Methods
Supplementary Text
Figs. S1 to S3
Tables S1 to S3
References

**Other Supplementary Materials for this manuscript include the following:**

Data S1 containing Tables S4 to S6

### Materials and Methods

Chemical Reagents

All chemical reagents were procured from external suppliers. Mezigdomide (CC-92480) was purchased as solid from SelleckChem catalogue number S8975, Iberdomide (CC-220) was purchased as solid from Stratech catalogue number S8760-SEL. Lenalidomide was purchased from Combi-Blocks catalogue number OR-2352. CFT-1297 was synthesized to order by Aragen Life Sciences.

Cloning

The stabilizing mutations in human CRBN (UniProt: Q96SW2) were designed using the PROSS server accessed in February 2022 (*1*). DNA encoding the CRBN constructs was synthesized as gBlock Gene Fragments by Integrated DNA Technologies and inserted into a pNIC28-Bsa4 (Addgene #26103) vector using ligation independent cloning, as well as into a pNIC28-Bsa4 with an additional maltose-binding protein (MBP) tag inserted between the $His_6$ and TEV (Tobacco Etch Virus) protease cleavage site. The final $CRBN^{midi}$ construct contained amino acids 41-187 followed by a Gly-Ser-Gly linker and amino acids 249-426 with single-point mutations C78I, I92V, K116N, Q134E, R283W, C287N, V293S, G302D, L342R, C343E, T359I and L423I.

Small-scale test protein expression

The $His_6$- and $His_6$-MBP-tagged CRBN fusion proteins were expressed in BL21(DE3) *E. coli* cells grown in 3 mL of Lysogeny broth (LB) supplemented with 50 µg/mL kanamycin. After 4.5 h at 37 °C, protein expression was induced with 0.5 mM isopropyl β-d-1-thiogalactopyranoside (IPTG) at 18 °C overnight. After harvesting, the cells were lysed in buffer containing 20 mM HEPES pH 8.0, 500 mM NaCl, 10 mM imidazole, 1 mM tris(2-carboxyethyl)phosphine hydrochloride (TCEP), 5 mM $MgCl_2$, cOmplete Protease Inhibitor (Roche), 0.5 mg/mL lysozyme, and 10 µg/mL DNase I (Roche) and incubated at RT for 30 mins. The lysate was clarified by centrifugation at 6,000 g for 20 mins at 4 °C and subsequently added to Ni-NTA Magnetic Agarose Beads (Qiagen). After extensive washing with buffer containing 20 mM HEPES pH 8.0, 500 mM NaCl, 20 mM imidazole, 1 mM TCEP, and 0.05% (v/v) Tween-20, the protein was eluted in buffer containing 20 mM HEPES pH 8.0, 500 mM NaCl, 250 mM imidazole, 1 mM TCEP, and 0.05% (v/v) Tween-20. The eluted protein was analyzed on NuPAGE 4-12% Bis-Tris polyacrylamide gels (Thermo Fisher Scientific) followed by InstantBlue Coomassie staining (Abcam).

$CRBN^{midi}$ expression and purification

The $His_6$-tagged fusion protein was expressed in BL21(DE3) *E. coli* cells grown in LB supplemented with 50 µg/mL kanamycin. Protein expression was induced with 0.5 mM IPTG at 18 °C overnight with 50 µM $ZnCl_2$ added at induction. After harvesting, the cells were resuspended in buffer containing 20 mM HEPES pH 8.0, 500 mM NaCl, 50 µM $ZnCl_2$, 0.5 mM TCEP, 0.05% (v/v) Tween-20. 5 mM imidazole, 1mM $MgCl_2$, cOmplete Protease Inhibitor (Roche), and 10 µg/mL DNase I and lysed using a Continuous Flow Cell Disruptor (Constant Systems) at 30,000 psi. The lysate was clarified by centrifugation at 48,000 g for 30 mins at 4 °C and loaded onto a HisTrap HP (Cytiva) column. Impurities were washed with buffer containing 20 mM HEPES pH 8.0, 500 mM NaCl, 0.5 mM TCEP, and 20 mM followed by 60 mM imidazole. The protein was eluted by increasing the imidazole concentration to 100-125 mM. The eluted protein was loaded onto a HiPrep Sephadex G-25 Desalting Column pre-equilibrated with 20 mM HEPES pH 8.0, 500 mM NaCl, 0.5 mM TCEP to remove imidazole. The protein was incubated overnight at 4 °C with $His_6$-tagged TEV protease. The TEV protease, uncleaved protein, $His_6$ tag, and remaining impurities were removed by reverse $Ni^{2+}$ affinity chromatography using HisTrap HP columns with $CRBN^{midi}$ eluting in buffer containing 10-20 mM imidazole, 20 mM HEPES pH 8.0, 500 mM NaCl, and 0.5 mM TCEP. The protein-containing fractions were concentrated (Amicon Ultra centrifugal filter, MWCO 10 kDa, Merck) and further purified by size-exclusion chromatography (SEC) on a Superdex 200 column (Cytiva) in buffer containing 20 mM HEPES pH 7.5, 500 mM NaCl, and 0.5 mM TCEP. The protein was concentrated to around 3 mg/mL, flash frozen in liquid nitrogen and stored at -80 °C until further use. All chromatography purification steps were performed using an Äkta Pure (Cytiva) system at 4 or 20 °C.

$CRBN^{TBD}$ expression and purification

The glutathione-S-transferase (GST) fusion protein was expressed in BL21(DE3) *E. coli* cells grown in LB supplemented with 50 µg/mL kanamycin. Protein expression was induced with 0.5 mM IPTG at 18 °C overnight with 50 µM $ZnCl_2$ added at induction. After harvesting, the cells were resuspended in buffer containing 50 mM HEPES pH 8.0, 500 mM NaCl, 1 mM TCEP, 1 mM $MgCl_2$, cOmplete Protease Inhibitor (Roche), and 10 µg/mL DNase I (Roche) and lysed using Continuous Flow Cell Disruptor (Constant Systems) at 30,000 psi. The lysate was clarified by centrifugation at 20,000 rpm for 40 mins at 4 °C and added to 7 mL of glutathione agarose beads equilibrated in 50 mM HEPES pH 8.0, 500 mM NaCl and 1 mM TCEP. The lysate was incubated for 2 hours at 4 °C before it was applied to a batch column and the impurities were washed with 50 mM HEPES pH 8.0, 500 mM NaCl and 1 mM TCEP. The protein was incubated overnight at 4 °C with $His_6$-tagged TEV protease for on-bead cleavage of the GST tag. The cleaved protein was eluted by washing the beads with 50 mM HEPES pH 8.0, 500 mM NaCl and 1 mM TCEP and collecting the flow through. The protein was concentrated (Amicon Ultra centrifugal filter, MWCO 3 kDa, Merck) and further purified by SEC on a Superdex 75 column (Cytiva) in buffer containing 20 mM HEPES pH 7.5, 200 mM NaCl, and 1 mM TCEP. The protein was concentrated to around 20 mg/mL, flash frozen in liquid nitrogen and stored at -80 °C until further use. All chromatography purification steps were performed using an Äkta Pure (Cytiva) system.

$BRD4^{BD2}$ expression and purification

Human $Brd4^{BD2}$ (residues 333-460, UniProt ID: O60885) was expressed with an N-terminal $His_6$ tag in *E. coli* BL21(DE3) at 37 °C with LB supplemented with 50 µg/mL kanamycin once $OD_{600}$ reached 0.8. Protein expression was induced with 0.5 mM IPTG with 50 µM $ZnCl_2$ added at induction and grown overnight at 18 °C. After harvesting, the cells were resuspended in buffer containing 12 mM phosphate buffer, 500 mM NaCl, 40 mM imidazole, 1 mM $MgCl_2$ and DNase I (10 µg/mL), and cells were lysed using a Continuous Flow Cell Disruptor (Constant Systems) at 30,000 psi. Cell lysates were clarified by centrifugation at 18,000 g for 30 minutes at 4 °C. The lysate was filtered and loaded to HisTrap FF affinity column (GE Healthcare) and eluted with 12 mM phosphate buffer, 500 mM NaCl and 500 mM imidazole). The final purified proteins were concentrated to around 9 mg/mL and stored in 20 mM HEPES, pH 7.5, 500 mM NaCl and 0.5 mM

TCEP at -80 °C until further use. All chromatography purification steps were performed using a BioRad NGC system at 4 °C.

IKZF1$^{ZF2}$ expression and purification

IKZF1$^{ZF2}$, which was incorporated into a pGEX6P-1 vector, was expressed in *E. coli* BL21(DE3) using LB supplemented with 50 μM $ZnSO_4$ and 100 μg/mL ampicillin. Protein expression was induced using 0.5 mM IPTG when $OD_{600}$ reached 0.6 - 0.8. The cells were harvested through centrifugation and subsequently re-suspended in a buffer comprising 20 mM Tris-HCl pH 8.0, 500 mM NaCl, 5% (v/v) glycerol, and 0.1 mM dithiothreitol (DTT). The protein purification was performed as previously described (*2*) with gel filtration conducted using a buffer composed of 50 mM HEPES pH 7.5, 150 mM NaCl, and 0.25 mM TCEP.

Crystallization of apo CRBN$^{midi}$

Crystals of apo CRBN$^{midi}$ were grown using the sitting drop vapour diffusion method at 20 °C by mixing equal volumes of protein solution (2.9 mg/mL in SEC buffer) with reservoir solution containing 0.2 M sodium citrate, 20% (w/v) PEG 3350, and 0.1 M BIS-TRIS propane pH 6.5. The crystal was cryoprotected in the reservoir solution supplemented with 20 % (v/v) glycerol and subsequently flash cooled in liquid nitrogen.
Diffraction data were collected at Diamond Light Source beamline I24 using a Pilatus3 6M detector at 100 K and 1.00 Å wavelength. The data was processed using autoPROC (*3*) and indexed in space group $C222_1$. The structure was solved by molecular replacement in phenix.phaser (*4*) using the atomic coordinates of the mezigdomide-bound CRBN$^{midi}$ structure as search model, assuming one molecule in the asymmetric unit. Refinement and model building was done iteratively using phenix.refine (*5*) and coot (*6*). The Ramachandran statistics for the final structure were: 97.5% favoured, 2.5% allowed and 0% outliers.

Crystallization of CRBN$^{midi}$ bound to mezigdomide

Crystals of CRBN$^{midi}$ bound to mezigdomide were grown using the sitting drop vapour diffusion method at 4 °C by mixing equal volumes of protein solution (4 mg/mL) supplemented with 214 μM mezigdomide in buffer containing 20 mM HEPES pH 7.5, 500 mM NaCl, 0.5 mM TCEP, 1% (v/v) DMSO with reservoir solution containing 0.2 M sodium acetate, 25% (w/v) PEG 3350, and 0.1 M BIS-TRIS pH 6.5. The crystals were cryoprotected in the reservoir solution supplemented with 20 % (v/v) ethylene glycol and subsequently flash cooled in liquid nitrogen.
Diffraction data were collected at Diamond Light Source beamline I24 using a Pilatus3 6M detector at 100 K and 1.00 Å wavelength. The data was processed using autoPROC (*3*) and STARANISO (*7*), indexed in space group $P4_32_12$. The structure was solved by molecular replacement in phenix.phaser (*4*) using a model predicted by ColabFold (*8*) for the CRBN$^{midi}$ sequence as a search model, assuming one molecule in the asymmetric unit. Refinement and model building was done iteratively using phenix.refine (*5*) and coot (*6*). The Ramachandran statistics for the final structure were: 95.8% favoured, 4.2% allowed and 0% outliers.

Crystallization of CRBN$^{midi}$ bound to lenalidomide

CRBN$^{midi}$ at 3 mg/mL in a buffer containing 20 mM HEPES pH 7.5, 500 mM NaCl and 0.5 mM TCEP was mixed with a 4-molar excess of lenalidomide (320 μM, 1.7% (v/v) DMSO final). The complex was subjected to co-crystallization using the sitting drop vapour diffusion method across several sparse matrix screens at 20 °C by mixing equal volumes of protein solution and reservoir

solution. Initial crystal hits from ProPlex well C5 (0.1 M Tris pH 8, 20% (w/v) PEG 4000) were optimized using the Hampton Research additive screen and the best diffracting crystals resulted from a reservoir solution containing 0.1 M Tris pH 8, 20% (w/v) PEG 4000 and 1.2% myo-inositol. The crystals were harvested directly from the drop and flashed cooled in liquid nitrogen.
Diffraction data were collected at Diamond Light Source beamline i24 using the CdTe Eiger2 9M detector at a wavelength of 0.6199 Å. The 2.5 Å dataset was processed using autoPROC (*3*), indexed in $C222_1$ space group and the structure was solved by molecular replacement in phenix.phaser (*4*) using the CRBN$^{midi}$:mezigdomide coordinates as a search model. Phenix.refine (*5*) and coot (*6*) were used for refinement and model building. The Ramachandran statistics for the final structure were: 93.29 % favored, 6.71 % allowed and 0 % outliers.

Crystallization of a CRBN$^{midi}$:CFT-1297:BRD4$^{BD2}$ ternary complex
CRBN$^{midi}$, CFT-1297 and Brd4$^{BD2}$ were mixed at a 1:4:4 ratio and purified as a ternary complex by SEC using a Superdex-75 10/300 Increase GL column (GE Healthcare). The ternary complex-containing fractions were pooled and concentrated to a final concentration of 3.5 mg/mL. The ternary complex was crystallized using the sitting-drop vapour diffusion method by mixing equal volumes of protein with 20% (w/v) PEG 3350, 0.2 M sodium malonate and 0.1 M Bis-Tris Propane pH 6.5. Small crystals which appeared after one week were used as a seeding stock to optimize the crystal size. The seed was diluted with 300 uL of the crystallization condition. The final crystals were grown by mixing the ternary complex, seed stock, and a mixture of 20% (w/v) PEG 3350, 0.2 M sodium citrate and 0.1 M Bis-Tris Propane pH 7.5 with a ratio of 1:0.1:0.9. Crystals appeared in one day and grew to their full size in one week at 20 °C. The crystals were harvested and flash-cooled in liquid nitrogen using 30% (v/v) glycerol in the crystallization solution as a cryo-protectant.
Diffraction data were collected at Diamond Light Source beamline I24 using the Eiger CdTe 9M detector at a wavelength of 0.6199 Å. Indexing and integration of reflections were performed using xia2.dials, and scaling and merging with xia2.multiplex (*9*). The space group was determined to be *P*1. The structure was solved by molecular replacement in phenix.phaser (*4*) using CRBN$^{midi}$:mezigdomide and Brd4$^{BD2}$ (PDB ID: 2OUO) as search models. Two instances of the ternary complex were found in the asymmetric unit, indicating a final solvent content of 50.09% as calculated from the Matthews coefficient. The initial model was refined iteratively using COOT(*6*) and phenix.refine(*5*). Ligand structures and restraints were generated using the PRODRG server (*10*). The Ramachandran statistics for the final structure were: 96.6% favoured, 3.4% allowed and 0% outliers.

Crystallization of a CRBN$^{midi}$:mezigdomide:IKZF1$^{ZF2}$ ternary complex
CRBN$^{midi}$, mezigdomide and IKZF1$^{ZF2}$ were mixed in a 1:4:4 ratio. The complex was crystallized using the sitting drop vapour diffusion method by mixing equal volumes of the ternary complex (107 μM) and reservoir solution containing 25% (w/v) PEG 3350, 0.2 M ammonium sulphate and 0.1 M HEPES pH 7.5. The crystal was cryoprotected in the reservoir solution containing 30% (v/v) ethylene glycol and flash cooled in liquid nitrogen.
Diffraction data collection was carried out at Diamond Light source beamline I24 using an Eiger CdTe 9M detector at a wavelength of 0.9537 Å. The data was processed using xia2.dials (*9*). The crystal belonged to the space group $P12_11$. The structure was solved by molecular replacement in phenix.phaser (*4*) using chain A from CRBN$^{midi}$:CFT-1297:BRD4$^{BD2}$ and chain L from the CRBN:DDB1:pomalidomide:IKZF$^{ZF1}$ complex (PDB ID: 6H0F) as search models. The initial

model was refined iteratively using COOT (*6*) and phenix.refine (*5*). The Ramachandran statistics for the final structure were: 94.3% favoured, 5.7% allowed and 0% outliers.

Analysis of crystal structures

The RMSD values for all structural alignments were calculated using the 'super' command over Cα atoms in PyMOL (version 4.6, Schrödinger). Structure figures were generated in PyMOL (version 4.6, Schrödinger).

SAXS

SAXS data were collected at beamline B21 at the Diamond Light Source in SEC-SAXS configuration (March 2023) (*11*). $CRBN^{midi}$ was concentrated to 6.5 mg/mL in 20 mM HEPES pH 7.5, 500 mM NaCl, 0.5 mM TCEP immediately before the experiment. Binary complexes were formed by addition of 400 µM of the compound in 4% DMSO and incubation on ice for 30 minutes. Samples were run on a Superdex S200 3.2/300 column (Cytiva) in 20 mM HEPES pH 7.5, 500 mM NaCl, 0.5 mM TCEP with 0.075 mL/min flowrate at 298 K coupled online to the SAXS beamline. SAXS data were acquired using a 3 second exposure at 12.4 kV and 3.6 m detector (Eiger 4M) distance across a q-range of $4.5\times10^{-3}$ - $3.4\times10^{-1}$ $Å^{-1}$. Frames from SEC-SAXS that corresponded to the main protein peak, were merged using Chromixs (*12*) and analyzed using the ATSAS (version 3.2.1) package (*13*). Radii of gyration and molecular mass were extrapolated from the Guinier plot using Primus. Pairwise-distance distributions were calculated using GNOM (*14*). The dimensionless Kratky plot (*15*) was obtained by manually plotting $(qR_g)^2I(q)/I_0$ against $qR_g$.

The scattering curves were compared to scattering curves back-calculated from the crystal structure of $CRBN^{midi}$:mezigdomide in closed conformation and a homology model of the open conformation using CRYSOL(*16*). The open conformation model was obtained by homology modelling using the SWISS-MODEL webserver (accessed April 18th 2023) (*17*) with the cryo-EM structure of apo-CRBN (PDB ID: 8CVP) (*18*) as the input structure.

Surface Plasmon Resonance

All SPR measurements were carried out on a Biacore™ 8K (Cytiva). $CRBN^{midi}$ was immobilized *via* a $His_6$ tag on a NiHC 1500M chip (XanTec) at 25 °C. The chip surface was prepared for ligand ($CRBN^{midi}$) capture as per manufacturer's instructions, flowing 350 mM EDTA, followed by buffer, then loaded using 5 mM $NiCl_2$ followed by buffer. For binary measurements $CRBN^{midi}$ was diluted to 350 nM and flowed at 10 µL/min for 420 seconds to capture a final bound response of 240 RUs. For binary measurements, 10 nM of $CRBN^{midi}$ was flowed at 10 µl/min for 420 seconds to capture a final bound response of approximately 10,000 RUs. The buffer for immobilization was 50 mM Tris pH 8.0, 150 mM NaCl, 0.25 mM TCEP, 0.005% (v/v) Tween-20 and 2% (v/v) DMSO.

Kinetic measurements were derived from a multicycle kinetics measurement, with a 5-point 3-fold dilution series of analyte CFT-1297 (5 µM to 20.6 nM). For ternary experiments the analyte was supplemented with 50 µM $BRD4^{BD2}$. Each concentration point was flowed independently at 50 µL/min with an association time of 150 s followed by 500 s dissociation. All measurements were carried out at 20 °C. The running buffer for these measurements was the same as for

Data was analyzed using Biacore$^{TM}$ Insight Evaluation Software (version 4.0.8.20368). Raw sensograms were solvent corrected, followed by reference and blank subtraction. To calculate $K_D$ values, data was globally fitted to 1:1 binding model. Alpha, a parameter indicative of cooperativity, was calculated by dividing binary $K_D$ by ternary $K_D$.

TR-FRET

A half-log dilution series of CFT-1297 was incubated with $His_6$-tagged CRBN constructs and Cy5-labelled BRD4$^{BD2}$ in TR-FRET buffer (50 mM HEPES pH 7.5, 150 mM NaCl, 0.5 mM TCEP, 0.01% (v/v) Tween-20, 2% (v/v) DMSO) for 30 minutes before addition of α-his Eu donor beads (20 ug/mL, Perking Elmer). The mixture was incubated in the dark with gentle rocking for 1 hour before readout using a PHERAstar microplate reader (BMG Labtech). All incubations were conducted at 20 ºC. Data were fitted by non-linear regression (curve fit) using Prism software (GraphPad version 10). Experiments were conducted in triplicate and were the average of three repeats.

Differential Scanning Fluorimetry

Samples were prepared in a total volume of 25 µL by mixing 4.8 µM CRBN$^{midi}$ or 3.2 µM CRBN$^{\Delta 40}$:DDB1$^{\Delta BPB}$ (purchased from Selvita) with SYPRO orange (Merck, S5692) at 5× final concentration and 100 µM lenalidomide when applicable in a final buffer containing 50 mM HEPES pH 7.5, 200 mM NaCl, 0.5 mM TCEP and 4% (v/v) DMSO. The temperature was increased from 25 to 95 °C at a rate of 1 °C/min using a CFX96 Real-Time C1000 Touch Thermal Cycler (Biorad) instrument. The melting temperature was calculated as the average of the minima of the derivative (-dRFU/d*T*) of two replicate samples. The melting curves were plotted using Prism software (GraphPad version 10).

Isothermal Titration Calorimetry

Experiments were carried out on an ITC200 instrument (Malvern) in buffer containing 50 mM HEPES pH 7.5, 500 mM NaCl, 1 mM TCEP, 4% (v/v) DMSO at 298 K stirring the sample at 600 rpm. The ITC titration consisted of a 0.4 µL initial injection (discarded during data analysis) followed by 19 × 2 µL injections with 180 s spacing between injections. Lenalidomide (400 µM) was directly titrated into CRBN$^{midi}$ (20 µM). For a control titration, 400 µM lenalidomide was titrated into buffer. The data was fitted using a one-set-of-site binding model to obtain dissociation constants, binding enthalpy (ΔH), and stoichiometry (N) using MicroCal PEAQ-ITC Analysis Software (version 1.1.0.1262, Malvern).

Molecular Dynamics (MD)

The CRBN$^{midi}$:mezigdomide crystal structure was pre-treated using the Protein Preparation Workflow (PPW) (*19*) available in Schrödinger Suite (Release 2022-2) before initiating the MD simulation experiments. The PPW treatment involved preparing the structure at pH 7.4, by adding hydrogen atoms, capping of the N and C termini with ACE (N-acetyl) and NMA (N-methyl amide) groups respectively, and building the missing side chain atoms. The missing residues were built using Prime (*20*). The hydrogen bond network was optimized by reorienting hydroxyl and thiol groups, water molecules, amide groups of asparagine (Asn) and glutamine (Gln), and the imidazole ring in histidine (His); and predicting protonation states of histidine, aspartic acid (Asp) and glutamic acid (Glu) and tautomeric states of histidine. The resulting structure was minimized to relieve any strain and fine-tune the placement of various groups. Hydrogen atoms were optimized fully, which allows relaxation of the H-bond network. Heavy atoms were restrained, so that a small amount of relaxation is allowed (RMSD = 0.30 Å). Mezigdomide-bound CRBN wild type structure as obtained from the Protein Data Bank (PDB ID: 8D7U) was also subjected to identical pre-treatment with the PPW. The final structures from the PPW were used as input for MD simulations. OPLS4 force field (*21*) was used throughout the computational studies.

MD simulations were carried out using the Desmond Molecular Dynamics System (*22*) implemented within the Schrödinger Suite (Release 2022-2). The systems were built with the input structures being placed in an orthorhombic box (with buffer distance of 15 Å) of explicit water molecules (TIP3P water model). A salt concentration of 0.15 M was maintained and ions ($Na^+$, $Cl^-$) were placed to neutralize the systems. The systems were minimized and then subjected to 100 ns of MD simulation runs with a recording interval of 10 ps for trajectory and energy. NPT ensemble class (pressure = 1 bar; temperature = 298.15 K) was used. RESPA integrator with 2 fs for bonded and near and 6 fs for far timesteps was used. Noose-Hoover chain thermostat (relaxation time = 1 ps) and Martyna-Tobias-Klein barostat (relaxation time = 2 ps) methods were used. A cut-off radius of 9 Å was set for coulombic interactions. For each system, five different simulations of 100 ns each were run with randoms seeds and randomizing the velocities at the beginning of the calculations. The Root Mean Square Fluctuations (RMSF) of protein residues as derived from the MD simulations and reported in this study are the averages over the five independent runs.

**Supplementary Text**

Molecular dynamics simulation

The RMSF of the protein residues in CRBN$^{\Delta N}$ and CRBN$^{midi}$ as observed during the MD simulation runs indicate that the two protein constructs exhibit a similar trend in their dynamic behavior (Fig. S2A,B). RMSF differences observed at a few localized regions of CRBN$^{midi}$ *vs.* CRBN$^{\Delta N}$ are far away from the ligand binding site and are mostly on the loops and/or solvent exposed (Refer to 'remarks' column of Table S4-S6). The median RMSF of the protein residues in Lon, HB and TBD domains of CRBN$^{midi}$ is lower than that in the CRBN$^{\Delta N}$ construct [Median RMSF in the Lon+HB domain: 1.24 Å (CRBN$^{\Delta N}$); 0.99 Å (CRBN$^{midi}$). Median RMSF in the TBD: 1.17 Å (CRBN$^{\Delta N}$); 0.94 Å (CRBN$^{midi}$)] (Fig. S2C,D). These observations indicate the mutations introduced in our CRBN$^{midi}$ construct perhaps impart rigidity and hence contribute towards stabilizing the protein. Further, the key interactions and their stabilities over the course of the MD simulations mediated by the glutarimide ring of mezigdomide with the aromatic cage of the protein remain preserved in CRBN$^{midi}$ as seen in CRBN$^{WT}$ (Fig. S2E). Overall, the computational investigations suggest that CRBN$^{midi}$ and CRBN$^{WT}$ show similar profiles for protein dynamics and ligand-mediated interactions with the TBD.

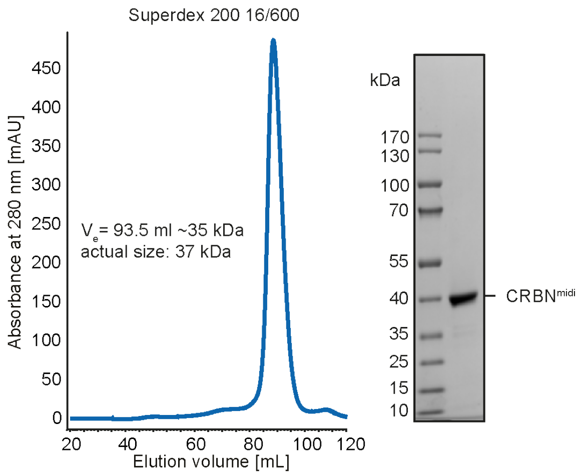

**Fig. S1. Purification of CRBN$^{midi}$.** SEC profile of purified CRBN$^{midi}$ (left) and SDS-PAGE analysis of the elution peak (right). The estimated size of the eluted protein calculated from a standard calibration curve is indicated.

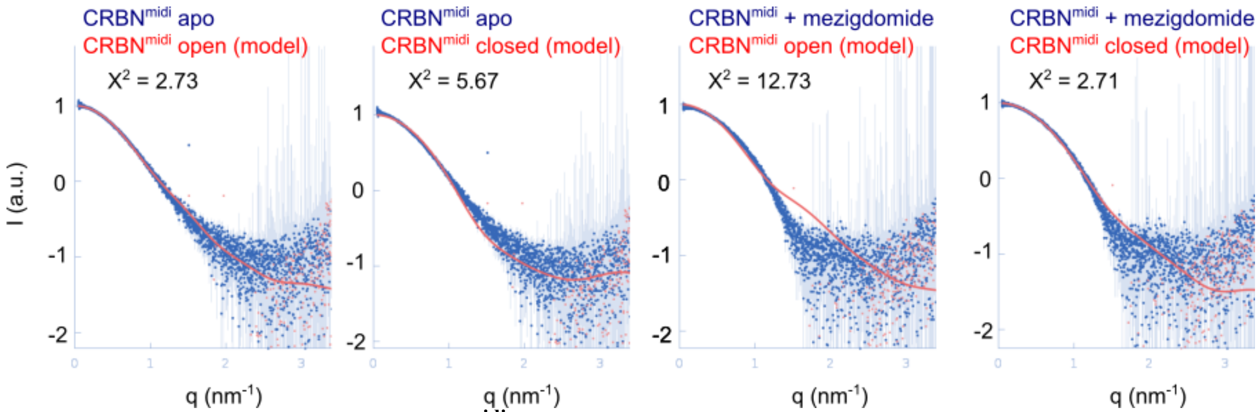

**Fig. S2. SAXS analysis of CRBN$^{midi}$ in the absence and presence of ligands.** Experimental (blue) scattering curves for apo and mezigdomide-bound CRBN$^{midi}$. Theoretical scattering curves (red) for open and closed CRBN$^{midi}$ models. The scattering curve for apo CRBN$^{midi}$ is in good agreement with the open model. In contrast, apo CRBN$^{midi}$ does not agree as well with the closed model. CRBN$^{midi}$:mezigdomide is likely in a closed conformation which is demonstrated by poor agreement with the theoretical open model curve and better agreement with the theoretical closed model curve.

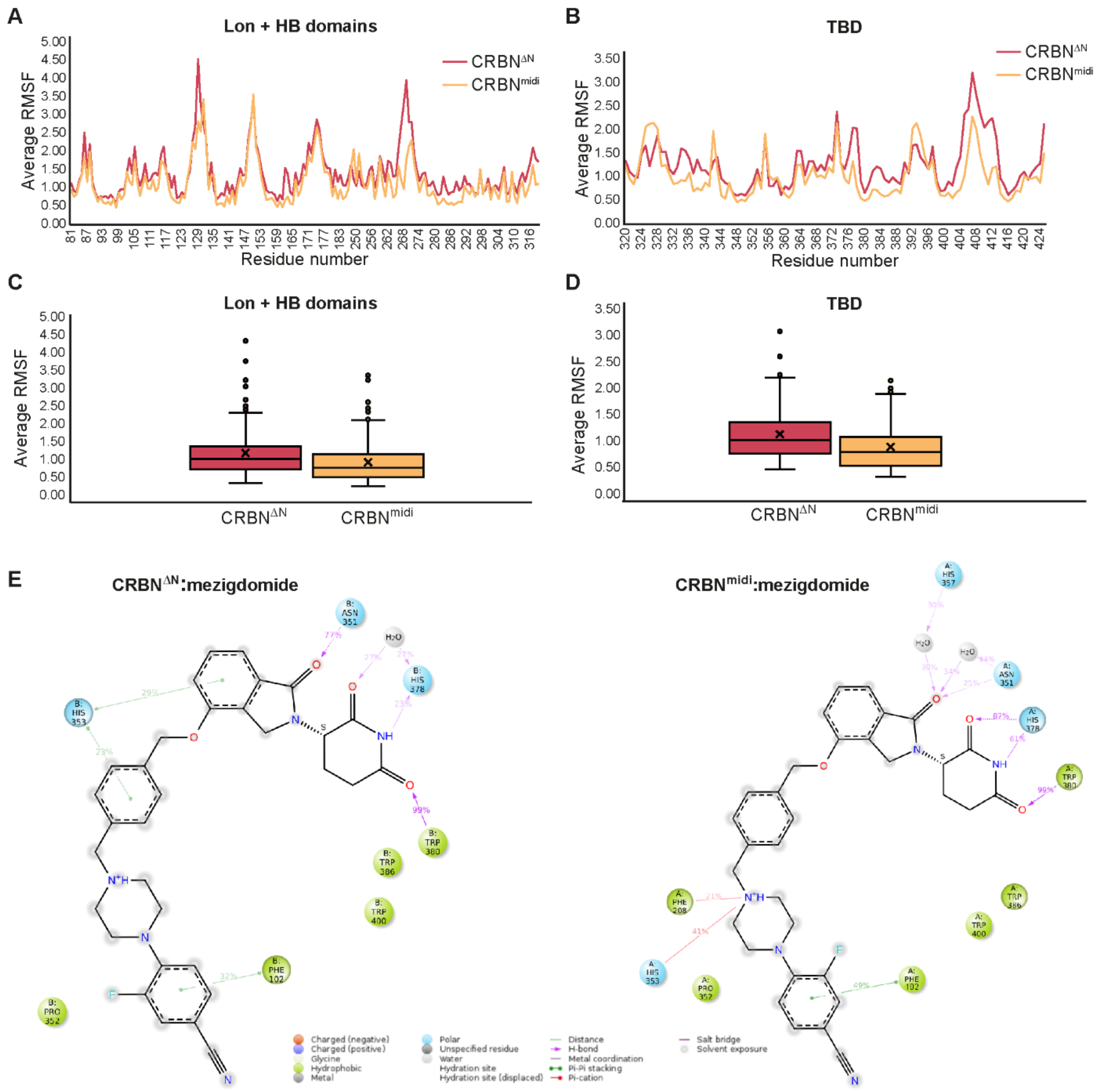

**Fig. S3. Molecular dynamics simulations. (A-D)** Average RMSF of the protein residues in CRBN$^{\Delta N}$ (red) *vs*. CRBN$^{midi}$ (orange). Residue-wise distribution of average RMSF of the protein residues in the Lon+HB domains (A) and the TBD (B). Box plots showing the distribution spread of average RMSF of the protein residues in the Lon+HB domains (C) and the TBD (D). **(E)** Protein-ligand interaction stability (in percentages) as observed in a representative MD simulation run of CRBN$^{\Delta N}$:mezigdomide (Left) and CRBN$^{midi}$:mezigdomide complexes (Right).

**Table S1. Data collection and refinement statistics for $CRBN^{midi}$ apo and binary structures.**

| | apo $CRBN^{midi}$ | $CRBN^{midi}$:mezigdomide | $CRBN^{midi}$:lenalidomide |
|---|---|---|---|
| **Data collection** | | | |
| Space group | $C222_1$ | $P4_32_12$ | $C222_1$ |
| Cell dimensions | | | |
| *a*, *b*, *c* (Å) | 52.03, 96.41, 148.09 | 51.05, 51.05, 267.68 | 51.99, 95.72, 148.25 |
| α, β, γ (°) | 90.0, 90.0, 90.0 | 90.0, 90.0, 90.0 | 90.0, 90.0, 90.0 |
| Resolution (Å) | 74.04 - 3.11 (3.16 - 3.11) | 66.92 - 2.19 (2.37 - 2.19) | 45.54- 2.5 (2.54 - 2.50) |
| $R_{merge}$ | 0.410 (6.797) | 0.244 (1.090) | 0.200 (1.782) |
| $I / \sigma I$ | 5.0 (0.7) | 6.4 (1.7) | 5.046 (1.17) |
| $CC_{1/2}$ | 0.98 (0.37) | 0.99 (0.53) | 0.99 (0.42) |
| Completeness (%) | 100.0 (100.0) | 91.8 (50.9) | 99.7 (96.4) |
| Redundancy | 11.7 (11.6) | 6.5 (4.1) | 4.8 (3.7) |
| **Refinement** | | | |
| Resolution (Å) | 74.04 - 3.11 | 40.59 - 2.19 | 45.54 - 2.5 |
| No. reflections | 6975 (688) | 13035 (653) | 13030 (1181) |
| $R_{work}$ / $R_{free}$ | 0.267 / 0.300 | 0.257 / 0.285 | 0.2994 / 0.342 |
| No. atoms | | | |
| Protein | 2119 | 2405 | 2267 |
| Ligand/ion | 1 | 43 | 20 |
| Water | 0 | 32 | 5 |
| *B*-factors | | | |
| Protein | 81.15 | 41.61 | 60.47 |
| Ligand/ion | 117.26 | 30.40 | 62.68 |
| Water | - | 41.61 | 39.78 |
| R.m.s. deviations | | | |
| Bond lengths (Å) | 0.003 | 0.009 | 0.015 |
| Bond angles (°) | 0.64 | 1.32 | 1.69 |
| **PDB ID** | 8RQ1 | 8RQ8 | 8RQA |

Each dataset was collected from a single crystal. Values in parentheses are for highest-resolution shell.

**Table S2. Data collection and refinement statistics for CRBN$^{midi}$ ternary structures.**

| | CRBN$^{midi}$:CFT-1297:BRD4$^{BD2}$ | CRBN$^{midi}$:mezigdomide:IKZF1$^{ZF2}$ |
|---|---|---|
| **Data collection** | | |
| Space group | *P*1 | *P*1211 |
| Cell dimensions | | |
| *a*, *b*, *c* (Å) | 43.88, 52.64, 130.35 | 53.56, 142.84, 56.69 |
| α, β, γ (°) | 96.44, 91.49, 99.22 | 90.00, 112.341, 90.00 |
| Resolution (Å) | 46.10 - 2.91 (2.96 - 2.91) | 52.44 - 2.15 (2.23 - 2.15) |
| $R_{merge}$ | 0.333 (3.092) | 0.200 (3.696) |
| $I / \sigma I$ | 3.3 (0.1) | 6.12 (0.32) |
| $CC_{1/2}$ | 0.99 (0.26) | 0.996 (0.343) |
| Completeness (%) | 97.8 (75.6) | 97.56 (80.10) |
| Redundancy | 6.6 (6.0) | 7.1 (7.2) |
| **Refinement** | | |
| Resolution (Å) | 43.27 - 2.91 | 52.44 - 2.15 |
| No. reflections | 20285 (275) | 41733 (3417) |
| $R_{work}$ / $R_{free}$ | 0.248 / 0.291 | 0.260 / 0.287 |
| No. atoms | | |
| Protein | 6236 | 5036 |
| Ligand/ion | 122 | 88 |
| Water | 31 | 206 |
| *B*-factors | | |
| Protein | 74.28 | 60.98 |
| Ligand/ion | 73.17 | 53.62 |
| Water | 54.08 | 63.50 |
| R.m.s. deviations | | |
| Bond lengths (Å) | 0.007 | 0.010 |
| Bond angles (°) | 1.13 | 1.45 |
| **PDB ID** | 8RQ9 | 8RQC |

Each dataset was collected from a single crystal. Values in parentheses are for highest-resolution shell.

**Table S3. CRBN$^{midi}$ immobilization level to Ni-NTA chip for SPR experiments and maximal response achieved by each analyte**

| Sample | CRBN$^{midi}$ immobilization level | Experimental $R_{max}$ | Theoretical $R_{max}$ | Experimental $R_{max}$ / Theoretical $R_{max}$ |
|---|---|---|---|---|
| **CFT1-297** | 9581 | 196 | 211 | 93% |
| **BRD4$^{BD2}$:CFT-1297** | 240 | 56 | 101 | 55% |

**Data S1. (separate file)**

Table S4. Comparison of average RMSF of protein residues in the topologically equivalent regions of the Lon domain of $CRBN^{\Delta N}$ *vs*. $CRBN^{midi}$ construct.

Table S5. Comparison of average RMSF of protein residues in the topologically equivalent regions of the TBD of $CRBN^{\Delta N}$ *vs*. $CRBN^{midi}$ construct.

Table S6. Comparison of average RMSF of protein residues at the mutation sites in $CRBN^{\Delta N}$ *vs*. $CRBN^{midi}$ construct.
